# Ultra-high-order ICA: fine overlapping functional parcellations and spatiotemporal reconfiguration

**DOI:** 10.1101/2020.03.11.986299

**Authors:** Armin Iraji, Zening Fu, Thomas DeRamus, Shile Qi, Srinivas Rachakonda, Yuhui Du, Vince Calhoun

## Abstract

Our recent findings show that functional organizations evolve spatially over time, highlighting the importance of considering within-subject spatial variations and dynamic functional parcellations in brain functional analyses. Meanwhile, a considerable level of multi-functionality suggests the need for overlapping brain parcellations. In this work, we used ultra-high-order ICA to identify fine overlapping functional dynamic parcellations of the brain. The preliminary result of this work was presented at the organization for human brain mapping workshop (OHBM 2019)^1^.

## Introduction

The brain reorganizes its activity interactively at different temporal and spatial scales. Recently, we demonstrated spatiotemporal reconfiguration of the brain’s functional organizations and changes within the region’s functional roles over time (A. Iraji, T. P. Deramus, et al., 2019; A. Iraji, Z. Fu, et al., 2019; A. Iraji, Miller, Adali, & Calhoun, 2020). Here, we present a framework using: 1) ultra-high-order independent component analysis (uICA) to parcellate the brain into dynamic fine overlapping functional units (FUs) (Armin Iraji et al., 2019) and 2) the concept of functional hierarchy to capture spatiotemporal functional reconfigurations of the brain. Our regularized cost function optimizes the selection of the best subsets of changes at each time-window and protects against spurious fluctuations.

## Methods

Spatial independent component analysis (sICA) has become an integral part of functional MRI (fMRI) studies, particularly with resting-state fMRI. Our recent work suggests that an ICA framework can identify granular and functionally homogeneous brain functional units (FUs). In this study, we applied uICA (1000 components) to estimate FUs at each time window. As a result, we obtained dynamic fine overlapping functional parcellation of the brain. Next, low-order ICA (20 components) was applied to identify large-scale intrinsic connectivity networks (LICNs), including the left and right frontoparietal, primary and secondary visual, somatomotor, default mode, cerebellar, salience, auditory, attention, and language networks. FUs were assigned to LICNs using the similarity of temporal activity between FUs and LICNs. First, the dominant (static) LICN assignment of each FU was obtained using the whole time series and all subjects. Next, for each time window (60 secs), FUs were assigned to LICNs using a novel regularized cost function (Figure 1). The cost function simultaneously maximizes the similarity between the temporal activities of a given FU, while minimizing the divergence from the dominant assignment. Therefore, it estimates FU assignments to LICNs while simultaneously protecting against the miscategorization of FUs due to spurious fluctuations and optimizing the selection of the best subsets of changes from the static group assignment.

**Figure 1.**
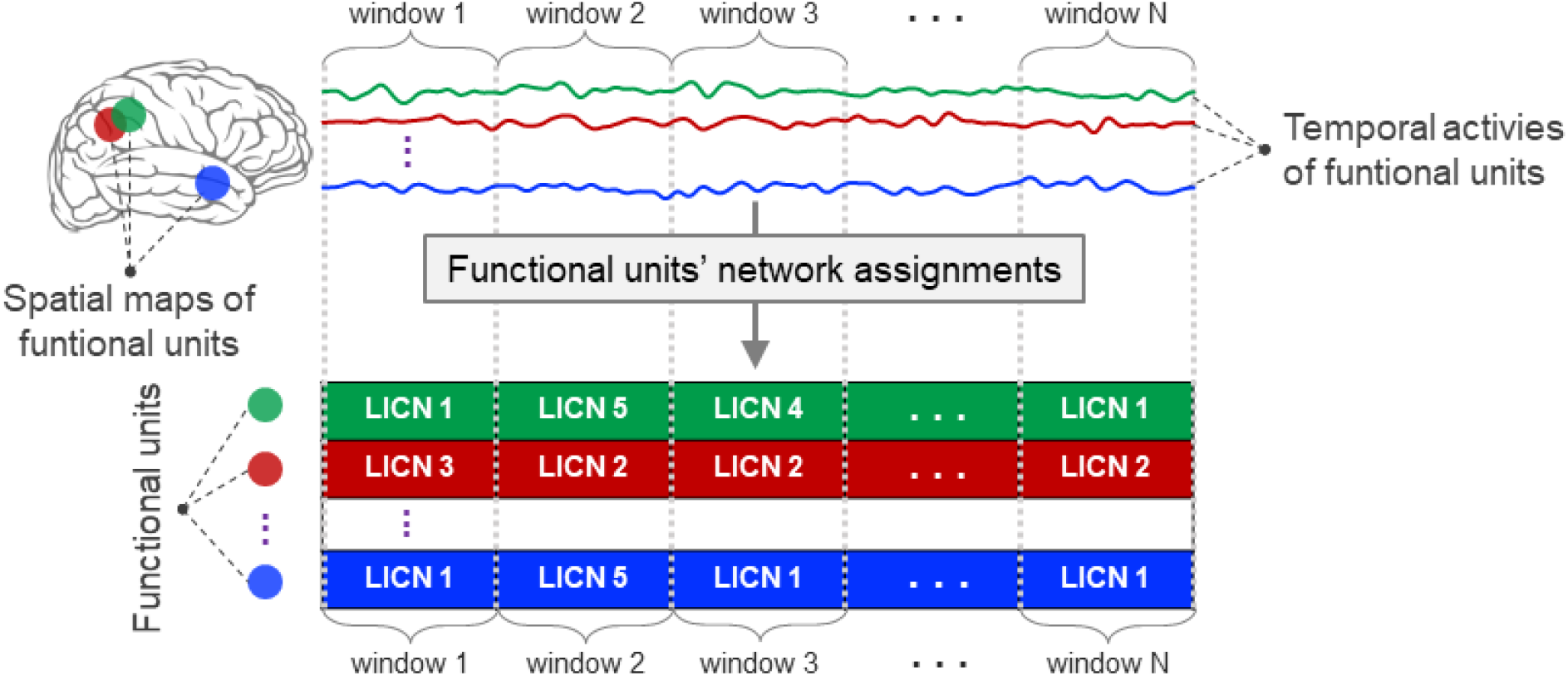
Evaluate the temporally dynamic configuration patterns of functional units (FUs) using the similarity of their temporal activities and the large-scale intrinsic connectivity networks (LICNs).

## Results

Figure 2(a) shows an example of brain parcellation designating primary FUs. The distribution of auxiliary FUs shows more than one FU are involved in every brain region, emphasizing the multi-functional roles of brain regions (Figure 2(b)). Spatially adjoined FUs reveal different functional roles over time, demonstrating the ability of uICA to identify fine distinct but also overlapping parcellations (FUs), in contrast to current parcellation approaches (example in Figure 2(c)). Further investigation shows reconfiguration in the spatial topography of LICNs over time. Spatial reconfiguration can be grouped into two categories: 1) expansion and contraction of networks spatial topography and 2) turnover in region membership. We also compared variation in FU assignments between-and within-subjects. Between-subjects variation is higher than within; however, the level of variations is within the same order of magnitude (Figure 2). Interestingly, results indicate a high level of agreement between variation across subjects and within subjects over time (correlation of 0.98). The network with higher (lower) variations across subjects, demonstrates lower (higher) within-subject variations. Furthermore consistent with previous findings using multi-task fMRI data, networks associated with higher cognitive functions have higher variation compared to networks previously found to be engaged in primary tasks (Cole et al., 2013; Dixon et al., 2018).

**Figure 2.**
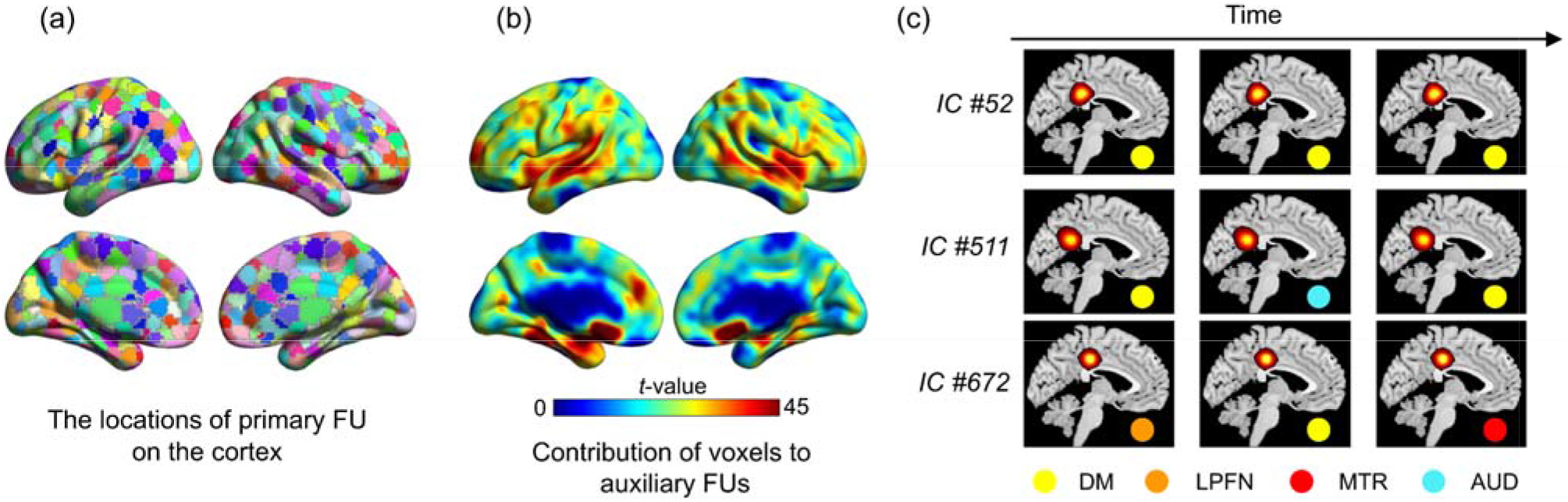
Spatial distribution of functional units (FUs) across the brain. The FUs were threshold at |Z| > 1.96 (p = 0.05). (a) The locations of the primary FUs on the cortex. Each voxel is assigned to the FUs with the maximum intensity. (b) Contribution of auxiliary FUs to each voxel. Brain regions can be involved in more than one FU. The colorbar represents the overall significant contribution of each voxel to FUs other than primary FU, emphasizing multiple roles of a given brain region to several brain functions. (c) An example of FUs obtained using ultra-high-order independent component analysis (uICA) and their assignment to large-scale intrinsic connectivity networks (LICN) as a function of time. UICA allows fine, overlapping parcellation of the brain using fMRI data. FUs demonstrate considerable variation in network assignments over time, and the variation is different even for overlapping FUs. Capturing overlapping FUs that have different patterns of activity and connectivity over time is one of the advantages of using uICA over current brain parcellation approaches. DM: default mode network; LPFN: left frontoparietal network; MTR: somatomotor network; and AUD: auditory network

**Figure 3.**
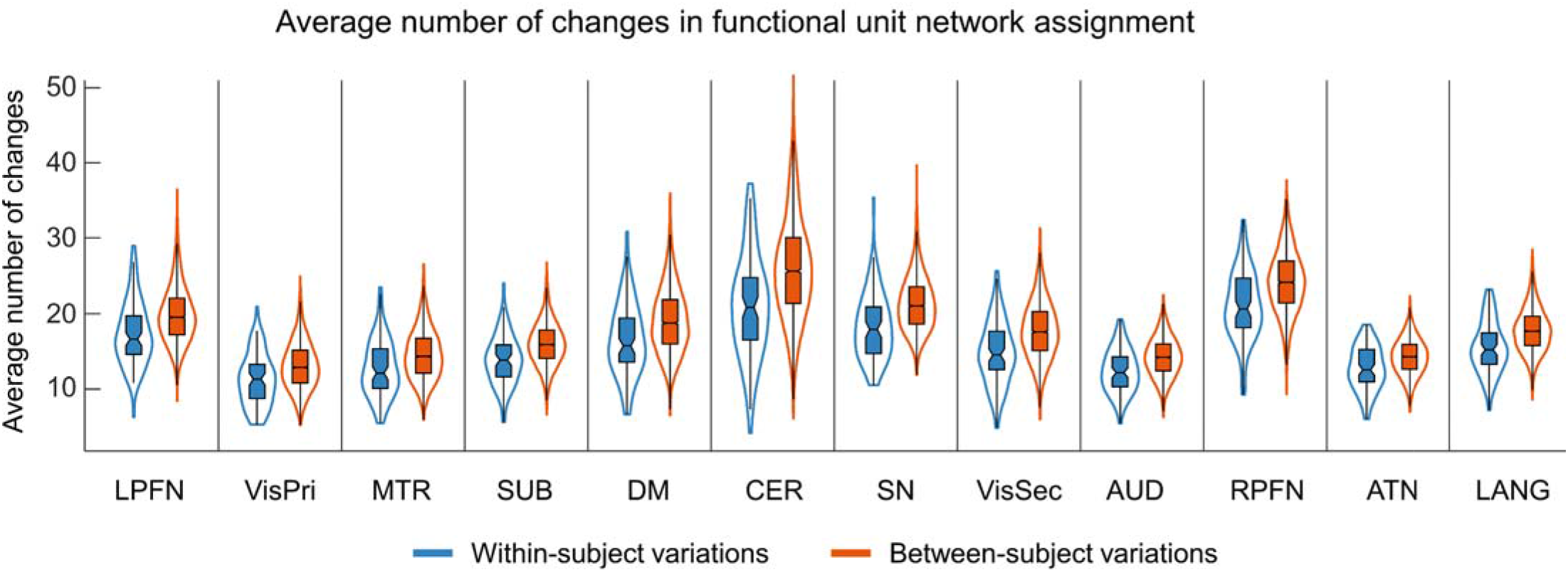
The average number of changes in functional units network assignments. Blue color indicates the within-subjects variations and orange color indicate between-subjects variations. LPFN: left frontoparietal network; VisPri: primary visual network; MTR: somatomotor network; SUB: subcortical; DM: default mode network; CER: cerebellar network; SN: salience network; VisSec: secondary visual network; AUD: auditory; RPFN: right frontoparietal; ATN: attention network; LANG: language network

## Discussion and Conclusion

We proposed a new data-driven (atlas-free) framework that captures spatiotemporal variation of brain function at a fine spatial scale and investigated the reconfiguration of functional, spatial topography (both between-and within-subjects). Preliminary results demonstrate the advantages of the proposed approach in investigating variation in brain functional organization. The findings show that changes in functional connectivity go beyond sampling variability, highlighting the importance of both between and within-subject variability. Temporal reconfiguration of brain spatial topography demands further investigation of the spatial chronnectome (Armin Iraji et al., 2018; A. Iraji et al., 2020).

## Footnotes

1 https://www.pathlms.com/ohbm/courses/12238/sections/15845/video_presentations/138027

## References

Cole, M. W., Reynolds, J. R., Power, J. D., Repovs, G., Anticevic, A., & Braver, T. S. (2013). Multi-task connectivity reveals flexible hubs for adaptive task control. Nat Neurosci, 16(9), 1348–1355. doi:10.1038/nn.3470

Dixon, M. L., De La Vega, A., Mills, C., Andrews-Hanna, J., Spreng, R. N., Cole, M. W., & Christoff, K. (2018). Heterogeneity within the frontoparietal control network and its relationship to the default and dorsal attention networks. Proc Natl Acad Sci U S A, 115(7), E1598–e1607. doi:10.1073/pnas.1715766115

Iraji, A., DeRamus, T., Lewis, N., Yaesoubi, M., Stephen, J. M., Erhardt, E., … Calhoun, V. (2018). The spatial chronnectome reveals a dynamic interplay between functional segregation and integration. bioRxiv, 427450. doi:10.1101/427450

Iraji, A., Deramus, T. P., Lewis, N., Yaesoubi, M., Stephen, J. M., Erhardt, E., … Calhoun, V. D. (2019). The spatial chronnectome reveals a dynamic interplay between functional segregation and integration. Hum Brain Mapp, 40(10), 3058–3077. doi:10.1002/hbm.24580

Iraji, A., Faghiri, A., Lewis, N., Fu, Z., DeRamus, T., Qi, S., … Calhoun, V. (2019). Ultra-high-order ICA: an exploration of highly resolved data-driven representation of intrinsic connectivity networks (sparse ICNs) (Vol. 11138): SPIE.

Iraji, A., Fu, Z., Damaraju, E., DeRamus, T. P., Lewis, N., Bustillo, J. R., … Calhoun, V. D. (2019). Spatial dynamics within and between brain functional domains: A hierarchical approach to study time-varying brain function. Hum Brain Mapp, 40(6), 1969–1986. doi:10.1002/hbm.24505

Iraji, A., Miller, R., Adali, T., & Calhoun, V. D. (2020). Space: A Missing Piece of the Dynamic Puzzle. Trends Cogn Sci, 24(2), 135–149. doi:10.1016/j.tics.2019.12.004

